# Antibody recognition of the Pneumovirus fusion protein trimer interface

**DOI:** 10.1101/2020.05.20.107508

**Authors:** Jiachen Huang, Darren Diaz, Jarrod J. Mousa

## Abstract

Human metapneumovirus is a leading cause of viral respiratory infection in children, and can cause severe lower respiratory infection in infants, the elderly, and immunocompromised patients. However, there remain no licensed vaccines or specific treatments for hMPV infection. Although the hMPV fusion (F) protein is the sole target of neutralizing antibodies, the immunological properties of hMPV F are still poorly understood. To further define the humoral immune response to the hMPV F protein, we isolated two new human monoclonal antibodies (mAbs), MPV458 and MPV465. Both mAbs are neutralizing *in vitro* and target a unique antigenic site harbored within the trimeric interface of the hMPV F protein. We determined both MPV458 and MPV465 have higher affinity for monomeric hMPV F than trimeric hMPV F. MPV458 was co-crystallized with hMPV F, and the mAb primarily interacts with an alpha helix on the F2 region of the hMPV F protein. Surprisingly, the major epitope for MPV458 lies within the trimeric interface of the hMPV F protein, suggesting significant breathing of the hMPV F protein must occur for hMPV F protein recognition of the novel epitope. In addition, significant glycan interactions were observed with a somatically mutated light chain framework residue. The data presented identifies a novel epitope on the hMPV F protein for structure-based vaccine design, and provides a new mechanism for human antibody neutralization of viral glycoproteins.

## Introduction

Human metapneumovirus (hMPV) is a leading cause of viral respiratory infections in children, the majority of which are seropositive for hMPV by five years of age^1^. Although hMPV was discovered in 2001^2^, there are no vaccines or therapeutics approved to prevent or treat viral infection. Similar to other respiratory pathogens, children, the elderly, and the immunocompromised are the major groups for which hMPV infection may require hospitalization^3–11^. Several reports have demonstrated hMPV infection can be lethal in both adults and children. In particular, haemopoietic stem cell transplant patients are at high risk of severe hMPV infection^10–13^, and several outbreaks of hMPV in nursing homes have been reported^14–16^. In addition, fatal hMPV has been observed in one child during an outbreak of hMPV in a daycare center.^17^ hMPV is also a significant cause of febrile respiratory illness in HIV-infected patients^18^, and has been linked to exacerbations of chronic obstructive pulmonary disease^19^. Co-circulation of hMPV was observed during the SARS outbreak of 2003, suggesting interactions with other circulating respiratory viruses.^20–22^

hMPV circulates as two genotypes, A and B, and based on the sequence variability of the surface proteins, hMPV is further grouped into four subgroups, A1, A2, B1, and B2^23,24^, and two additional subgroups, A2a and A2b, have been proposed^12^. hMPV has three surface glycoproteins, the small hydrophobic (SH), the attachment (G), and the fusion (F) proteins. The hMPV SH protein has been demonstrated to have viroporin activity^25^, while the hMPV G protein is thought to be involved in cellular attachment^26^. The hMPV F protein is indispensable for hMPV infection, and is highly conserved among hMPV subgroups^27^. Furthermore, the hMPV F protein is the sole target of neutralizing antibodies^28^. While the RSV G protein is immunogenic and elicits neutralizing antibodies^29^, the hMPV G protein is immunogenic, yet hMPV G-specific antibodies are non-neutralizing^1^. Although the hMPV G protein is thought to interact with proteoglycans, the hMPV F protein can interact with glycans in the absence of hMPV G.^30^ The hMPV F protein contains a highly conserved RGD motif that has been proposed as a key region in receptor binding to cellular integrins.^31,32^ The entry mechanisms of hMPV into the host cell membrane can occur by cell membrane or endosomal membrane fusion^33^.

Both hMPV and the related respiratory syncytial virus (RSV) share the *Pneumoviridae* family, and a similar F protein that has approximately 30% homology between the two viruses. For both viruses, the F protein has two long-lived conformations, the pre-fusion and post-fusion states^34^. Both RSV and hMPV lacking the G protein can infect cells *in vitro*, although these viruses are attenuated *in vivo*^35^. The pre-fusion conformation of the F protein is meta-stable, and stabilized versions of both hMPV F^36^ and RSV F^37,38^ have been generated. The RSV F protein was initially stabilized in the pre-fusion conformation using cysteine substitutions to lock the protein in the pre-fusion state by disulfide bonds, and through cavity-filling mutations to prevent transition to the post-fusion state. This Ds-Cav1 construct has been developed for clinical trials has shown promise in a phase I clinical trial^39^. Additional constructs for RSV F have focused on stabilizing the α4–α5 loop through proline mutations^38^. A similar approach was undertaken for the hMPV F protein, whereby a S185P mutation was introduced to stabilize the pre-fusion conformation^36^. The hMPV F protein contains a single site that is cleaved to convert the polypeptide F_0_ protein into the meta-stable disulfide-linked F_1_-F_2_ pre-fusion homotrimer. This is in contrast to RSV F, which contains two furin cleavage sites flanking the p27 peptide fragment. The cleavage enzyme for hMPV F *in vivo* is currently unknown, although cleavage can be accomplished by trypsin *in vitro*^40^. Post-fusion hMPV F was generated by removing the fusion peptide and incorporating one furin cleavage site from RSV F^41^. Based on these stabilized pre-fusion and post-fusion hMPV F constructs, X-ray crystal structures of the hMPV F protein from the A1 subgroup have been determined in the pre-fusion and post-fusion conformations^36,41^. Both proteins were expressed in CV-1 cells using a vaccinia virus expression system, although stabilized versions for routine HEK293F or CHO cell line expression have not yet been generated.

For RSV F, the pre-fusion conformation contains antigenic sites Ø^42^ and V^43^ located on the head of the F protein, which elicit the most potent neutralizing antibodies as compared to the post-fusion conformation^42,43^. Furthermore, the human antibody response to RSV infection is primarily focused on these pre-fusion-specific epitopes^44^. For hMPV F, data using human serum has shown that the preponderance of hMPV F-specific human antibodies bind both pre-fusion and post-fusion F conformations, which has been proposed is due to differential glycan positioning on the head of the hMPV F protein as compared to the RSV F protein^36^. Although several monoclonal antibodies (mAbs) have previously been isolated that recognize the hMPV F protein^41,45–52^, the predominant antigenic sites targeted by the human antibody response are unclear. A panel of rodent-derived mAbs was initially used to map the neutralizing epitopes on the hMPV F protein using viral escape mutants^45,46^. The known antigenic sites on the hMPV F protein include antigenic sites III, IV, and an unnamed site targeted by mAb DS7^34^. DS7 was isolated from a human phage display library^47^, and was co-crystallized with a fragment of the pre-fusion hMPV F protein^53^. Several mAbs isolated have been found to cross-neutralize RSV and hMPV, including MPE8^49^ and 25P13^50^ (site III), and 101F^41^, 54G10^48^, and 17E10^51^ (site IV). In addition, we have recently isolated a panel of human mAbs targeting site III and the DS7 epitope^52^. One of these mAbs, MPV364, competes for binding at antigenic site III, but does not cross-react with RSV F, suggesting further examination of hMPV F epitopes is required. In this study, we isolated new human mAbs to further identify the epitopes on the hMPV F protein recognized by the human immune system.

## Results

### Isolation of human antibodies to the hMPV F protein

To further identify the major antigenic epitopes on the hMPV F protein, we isolated mAbs from human subjects using hybridoma technology^54^. As hMPV infection and exposure is not routinely tested in patients, and the majority of individuals are seropositive for hMPV infection^55^, we isolated mAbs from two healthy human subjects. Two new mAbs were isolated against the recombinantly expressed hMPV B2 F protein (**Table S1**) expressed in HEK293F cells^52^. MPV458 and MPV465 were isolated from two different donors, and have isotypes of IgG_3_ and kappa, and IgG_1_ and lambda, respectively. MPV458 utilizes V_H_3-30, J_H_3, D_H_2, V_K_1-33, and J_K_5, while MPV465 utilizes V_H_3-33, J_H_5, D_H_3-22, V_L_-47, and J_L_3. The heavy chain complementarity determining region (HCDR) 3 loop length differs dramatically between the two mAbs as the HCDR3 loop for MPV458 is just eight amino acids, while MPV465 has a 21 amino acid long CDR3 loop (**Table S2**).

### Epitope identification

To identify the antigenic epitopes targeted by the isolated mAbs, we performed epitope binning using competitive biolayer interferometry^56^. Previously discovered mAbs with known antigenic epitopes were utilized as mapping controls, including mAbs 101F^57^ (site IV), MPV196^52^ and DS7^47^ (DS7 epitope), and MPE8^49^ and MPV364^52^ (site III) (**Fig. 1A**). Anti-penta-HIS biosensors were loaded with the hMPV 130-BV F^36^ protein and then loaded with one hMPV F-specific mAb, followed by exposure to a second mAb. mAbs MPV458 and MPV465 did not compete with any of the mapping control mAbs, yet competed for binding with each other, suggesting these two mAbs bind to a unique antigenic site on the hMPV F protein.

**Figure 1.**
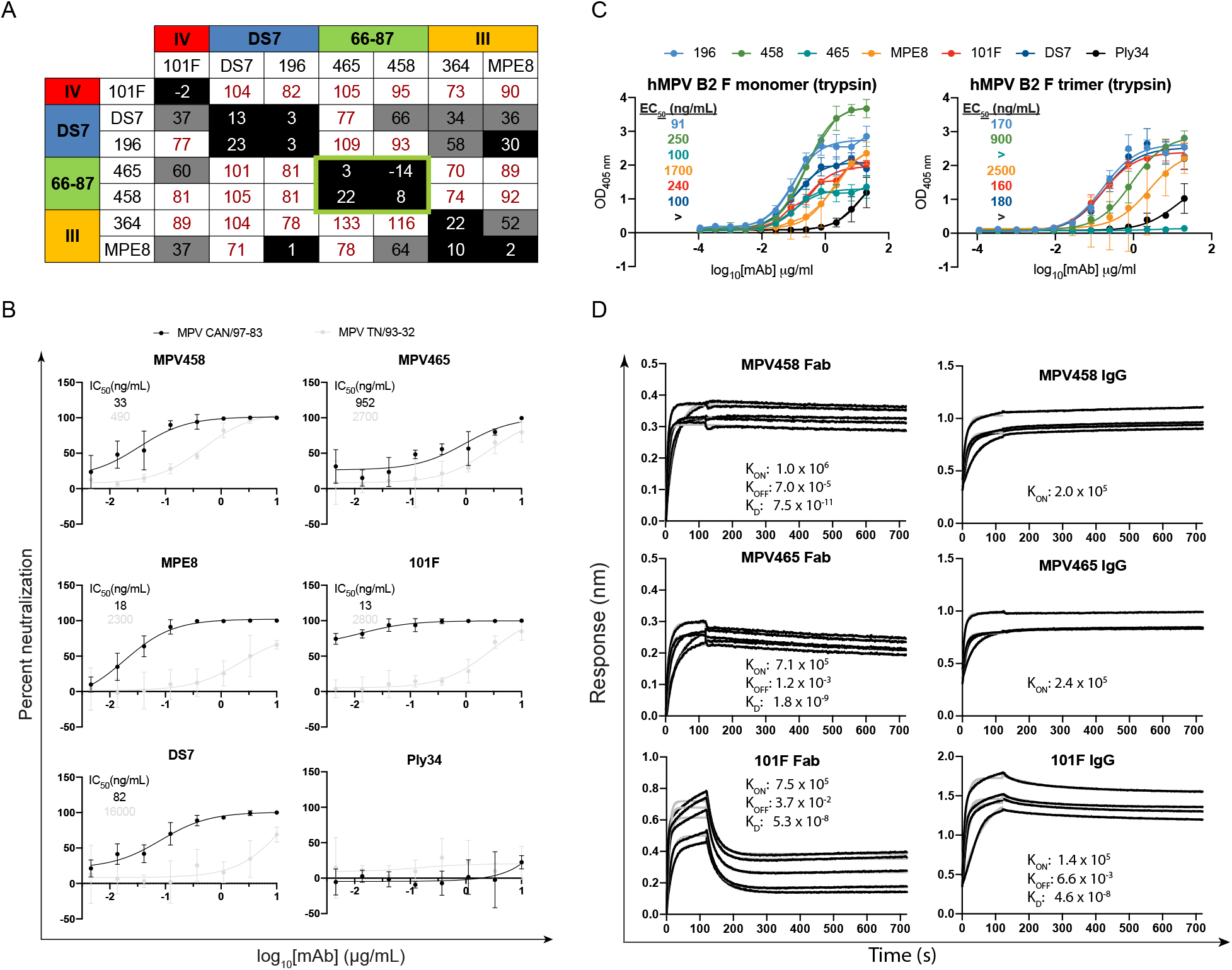
Binding and neutralizing properties of MPV458 and MPV465. (A) Epitope binning of the hMPV F-specific mAb panel. Epitope control mAbs include 101F (site IV), DS7 and MPV196 (DS7 epitope), and MPV364 and MPE8 (site III). MPV465 and MPV458 do not compete with known mAbs, and compete with each other for binding, suggesting both mAbs bind at a previously undiscovered antigenic site. Data indicate the percent binding of the competing antibody in the presence of the primary antibody, compared with the competing antibody alone. Cells filled in black indicate full competition, in which ≤33% of the uncompeted signal was observed; cells in gray indicate intermediate competition, in which the signal was between 33% and 66%; and cells in white indicate noncompetition, where the signal was ≥66%. Antigenic sites are highlighted at the top and side based on competition binding with the control mAbs. (B) Plaque neutralization curves for MPV458 and MPV465 with controls. Both MPV458 and MPV465 are neutralizing, while MPV458 has neutralizing properties similar to MPE8 and 101F. IC_50_ values are inlaid in each curve. The pneumococcal-specific antibody Ply34 was used as a negative control. Data points are the average of three replicates and error bars are 95% confidence intervals. Data are shown from one experiment and are representative of two independent experiments. A mAb was considered neutralizing if >50% plaque reduction was observed at the highest concentration of 20 μg/mL. (C) ELISA binding curves for hMPV F-specific mAbs against monomeric and trimeric hMPV B2 F protein that was treated with trypsin. MPV458 and MPV465 have lower EC_50_ values (higher affinity) for monomeric hMPV B2 F than trimeric hMPV B2 F. Binding curves and EC_50_ values are colored according to the legend. Each data point is the average of four replicates and error bars represent 95% confidence intervals. Data are representative of one experiment from two independent experiments. (D) Binding curves from biolayer interferometry. hMPV 130-BV coated anti-penta-HIS biosensors were exposed to mAbs for 120 s before dissociating in buffer for 600 s. Binding constants are displayed within each graph. For mAbs that exhibited limited dissociation, only association constants are displayed.

### Neutralization and binding properties

Plaque neutralization assays were performed to determine the neutralization properties of MPV458 and MPV465 against hMPV subgroup B2 (strain TN/93-32) and hMPV subgroup A2 (strain CAN/97-83) *in vitro* (**Fig. 1B**). MPV458 neutralized hMPV with 50% inhibitory concentration (IC_50_) values of 33 ng/mL for MPV CAN/97-83 and 490 ng/mL for MPV TN/93-32, while MPV465 had IC_50_ values of 950 and 2700 ng/mL, respectively. The neutralization potency of MPV458 was comparable to mAbs MPE8 and 101F. We next determined the binding properties of MPV458 and MPV465 by ELISA and biolayer interferometry. For ELISA, the half-maximal effective concentration (EC_50_) values were used to quantify binding between mAbs across multiple hMPV F protein constructs (**Table S1**, **Fig. S1-S4**). Generating trimeric hMPV F can be achieved by treating purified protein with trypsin as previously described^41,52^, although this process generates batch to batch variation of both pre-fusion and post-fusion conformations^52^. Both mAbs bind to hMPV F proteins from all four hMPV F subgroups (**Fig. S5**). We also quantified binding to hMPV F constructs that were predominantly in the pre-fusion and post-fusion conformations (**Fig. S5**). No major differences were observed between the predominantly pre-fusion hMPV F 130-BV protein and the predominantly post-fusion hMPV B2 GCN4 6R F protein, indicating these mAbs bind both pre-fusion and post-fusion conformations. We next assessed binding to exclusively monomeric and trimeric hMPV B2 F proteins that were treated with trypsin to induce cleavage (**Fig. 1C**). Both mAbs MPV458 and MPV465 had stronger binding to monomeric hMPV F than to trimeric hMPV F. MPV458 had a nearly four-fold lower EC_50_ to monomeric hMPV B2 F than to trimeric hMPV B2 F. MPV465 bound well to the hMPV B2 F monomer, while binding was completely abrogated binding to the hMPV B2 F trimer. These data indicate the epitope for MPV458 and MPV465 is predominantly exposed on monomeric hMPV F. Binding avidity and affinity were assessed by biolayer interferometry using the predominantly pre-fusion hMPV 130-BV protein (**Fig. 1D**). Affinity measurements were completed by cleaving mAbs to Fab fragments. MPV458 Fab had a faster K_ON_ than MPV465 Fab and 101F Fab, and also had limited dissociation, which gave a K_D_ 2-logs higher than MPV465 and 3-logs higher than 101F. Limited dissociation was observed for MPV458 and MPV465 IgG molecules as compared to 101F IgG, and thus a K_OFF_ rate could not be obtained. Overall, these data indicate MPV458 has higher affinity for the hMPV 130-BV F protein than mAbs MPV465 and 101F.

### X-ray crystal structure of the hMPV B2 F + MPV458 complex

To fully define the epitope targeted by the newly isolated mAbs, we co-crystallized the Fab of MPV458 in complex with hMPV B2 F. Trypsinization of hMPV B2 F generated trimeric and monomeric versions of hMPV F as assessed by size exclusion chromatography (**Fig. S6**). Cleavage of MPV458 and MPV465 mAbs to Fab fragments and subsequent addition of these Fabs to trypsinized trimeric hMPV B2 F resulted in monomeric hMPV F-Fab complexes (**Fig. S6**). Although the hMPV B2 F trimer appeared to fall apart upon Fab binding, we cannot attribute this to binding of MPV458 and MPV465 as other Fabs also caused trimer dissociation of this construct. The MPV458-hMPV B2 F complex was subjected to crystallization screening and crystals were obtained in 0.5 M ammonium sulfate, 0.1 M sodium citrate tribasic dihydrate pH 5.6, and 1.0 M Lithium sulfate monohydrate. Crystals were harvested and X-ray diffraction data was collected, and the structure of the complex was determined to 3.1 Å (**Fig 2**, **Table S3**). The asymmetric unit contained one hMPV F protomer with one MPV458-Fab molecule. hMPV F was observed in the pre-fusion conformation, although no trimeric structure was observed when viewing symmetry related partners (**Fig. S7**). MPV458 targets a unique epitope compared to previously discovered Pneumovirus antigenic sites. The primary epitope consists of a single alpha helix of amino acids 66-87 of the F2 region (**Fig. 2A**). Compared to the hMPV F protein, MPV458 binds nearly perpendicular to the long axis of the F protein. Upon overlay with the previously determined X-ray crystal structure of pre-fusion hMPV F, it is clear the major epitope lies completely within the interface between two protomers of trimeric hMPV F (**Fig. 2B**). This unusual epitope suggests the hMPV F protein is partially monomeric on the surface of the virion envelope or on virally infected cells. Alternatively, substantial breathing of the hMPV F protein could take place to allow the antibody to bind and neutralize the virus. As mentioned earlier, MPV458 has an unusually short HCDR3 loop of just 8 amino acids. The HCDR3 and light chain CDR (LCDR) 3 are centered on the 66-87 helix region. Numerous hydrogen bonding events were clear in the electron density (**Fig. 2C, 2D**, **S8**). The HCDR3 interacts via Asp107 with Arg79 of hMPV F, while HCDR2 Asn64 and Ser63 interact with Glu80 and Arg205, respectively (**Fig. 2C**). The HCDR1 utilizes Arg36 to interact with Glu70. The light chain LCDR3 has more hydrogen bonding events than the HCDR3, utilizing the backbone amino group of Leu114 to interact with Thr83, Arg115 hydrogen bonds to Asp87, and Asp108 bonds to Lys82 (**Fig. 2D**). LCDR1 Arg37 interacts with Asn57, which has an extended N-linked glycan motif. The LCDR2 Asp56 interacts with Thr56. The Framework 3 loop of the light chain interacts with the glycan motif consisting of NAG-NAG-BMA with branched MAN residues off the BMA glycan, in which Tyr83 interacts with the extended MAN glycan, while the long-face of Tyr83 site parallel to the extended glycan, suggesting a favorable interaction with the glycan motif.

**Figure 2.**
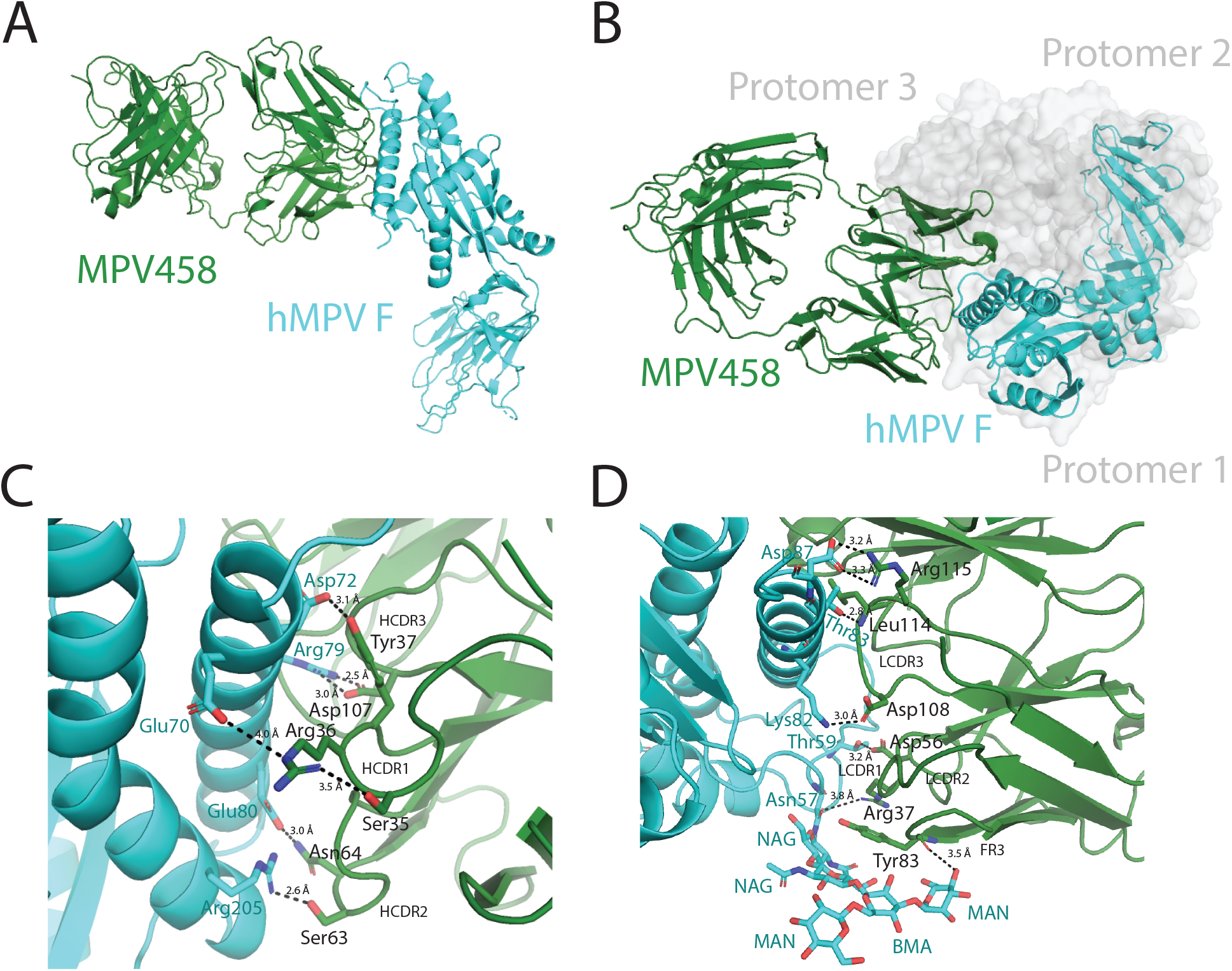
X-ray crystal structure of the hMPV B2 F + MPV458 Fab complex. (A) The asymmetric unit of the complex is displayed. Monomeric hMPV B2 F co-crystallized with one Fab of MPV458. (B) Overlay of the hMPV B2 F + MPV458 Fab complex with the previously determined X-ray crystal structure of hMPV A1 F in the pre-fusion conformation (MPV 115-BV, PDB: 5WB0). The hMPV F protein from each structure were overlaid in PyMol. MPV458 clashes with the trimeric structure. (C) Hydrogen bonding events observed between hMPV B2 F and the MPV458 Fab heavy chain. (D) Hydrogen bonding events observed between hMPV B2 F and the MPV458 light chain. The MPV458 light chain also interacts with an extended glycan patch linked from Asn57. CDR is complementarity determining region, FR is framework region. MPV458 numbering is in IMGT format.

### Functional characterization of the 66-87 helical epitope

The 66-87 helix of hMPV F is structurally conserved in the pre-fusion and post-fusion conformations, although the helix is exposed on the outer surface in the trimeric post-fusion conformation (**Fig. 3A, 3B**). Upon overlay of the 66-87 region of the pre-fusion and post-fusion hMPV F proteins, residues 66-83 align well, while the helix breaks on post-fusion hMPV F at residues 84-87 (**Fig. 3C)**. This sequence identity of the helix is highly conserved, as residues are identical between the A1 and B2 subgroups, except for a Lys82/Arg82 mutation. As MPV458 and MPV465 exhibited binding to post-fusion hMPV F constructs, we further examined binding by attempting to generate a complex between the Fab of MPV458 and trypsinized hMPV B2 F that was in the post-fusion conformation (**Fig. S2**, **S9**). No complex was observed as assessed by size exclusion chromatography while the Fab of 101F formed a complex with the post-fusion hMPV F protein. This suggests that although binding is observed by ELISA, the complete epitope lies outside the 66-87 helix and is incomplete in the post-fusion conformation. Since the major epitope is focused on the single helix, we assessed binding by Western blot to determine if MPV458 displayed binding to a linear conformation in the denatured hMPV F protein (**Fig. S10**). Binding to hMPV B2 F was analyzed using reduced and heated protein, and a nonreduced protein. MPV458 showed binding to all states of hMPV B2 F, while control mAbs 101F and MPE8 showed binding to only the nonreduced state. These data suggest the MPV458 epitope is at least partially linear. As the epitope for MPV458 lies within the trimer interface, the mechanism by which B cells recognize this epitope is unclear. To determine if the MPV458 epitope is exposed on the surface of virally infected cells, we performed flow cytometry using MPV458, MPE8, and a negative control pneumococcal-specific antibody (**Fig. S11**). Both MPV458 and MPE8 induced a fluorescent shift in virally infected cells, while the negative control mAb did not. This indicates the MPV F protein is either in monomeric form on the surface of infected cells, or that hMPV F trimer exhibits breathing motion that allows for binding of MPV458. By comparing the binding sites with previously described hMPV F-specific mAbs that have been structurally characterized (MPE8, 101F, DS7), the MPV458 epitope is distant from all three known antigenic sites (IV, VI, and III), and lies on the opposite face of the monomeric hMPV F protein (**Fig. 3D, 3E**). This unique epitope was unexpected on the hMPV F protein, although one intratrimeric epitope has recently been observed on the influenza hemagglutinin protein by mAb FluA-20^58^. However, FluA-20 was nonneutralizing and functioned by disrupting the HA trimer and inhibiting cell-to-cell spread. Evidence for Pneumovirus F protein breathing was previously demonstrated on the RSV F protein, whereby the mAb CR9501 that binds at antigenic site V enhances opening of the pre-fusion RSV F protein^59^.

**Figure 3.**
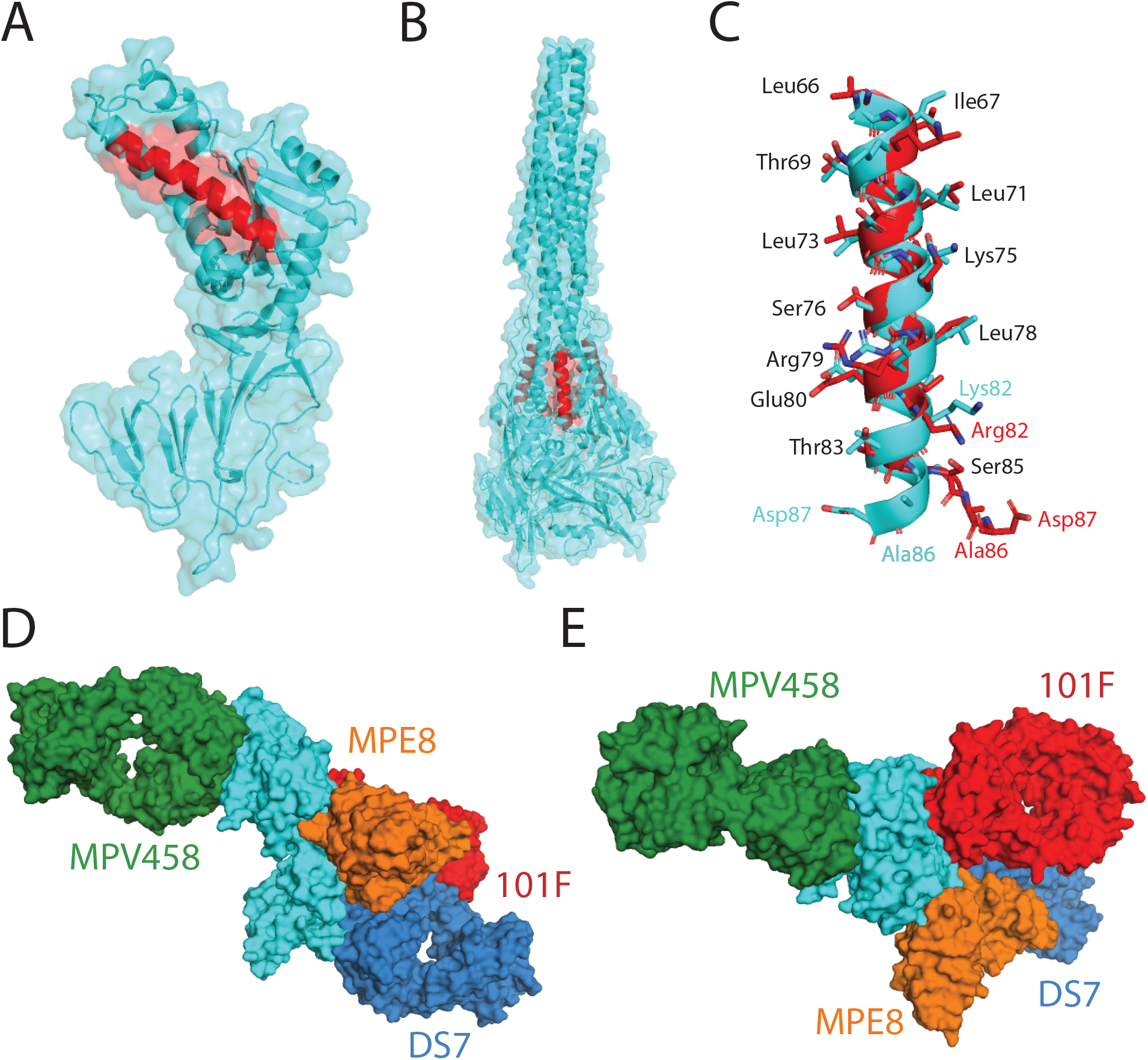
Structural comparison of the hMPV B2 F + MPV458 Fab complex. (A) The X-ray crystal structure of pre-fusion hMPV F is shown with the 66-87 epitope colored in red (PDB ID: 5WB0). (B) The corresponding 66-87 epitope is colored on the X-ray crystal structure of post-fusion hMPV F (5L1X). The 66-87 epitope is surface exposed on trimeric post-fusion hMPV F. (C) Structural overlay of the 66-87 region between pre-fusion (cyan) hMPV F from the hMPV B2 F + MPV458 Fab complex and post-fusion (red) hMPV F (PDB ID: 5L1X). Conserved amino acid residues between the B2 and A1 subgroups are listed in black, while residues that have mutations or shift positions are colored according to the corresponding structure. (D) Structural overlay of MPV458 on the hMPV F protein with previously structurally characterized hMPV F-specific mAbs. MPE8 (site III, orange) and 101F (site IV, red) were aligned onto hMPV F by aligning the corresponding RSV F residues onto hMPV F from the co-complex structures with RSV F (PDB ID: 5U68 and PBD ID: 3O45). DS7 was aligned from PDB 4DAG. (E) The structure overlay in (D) is rotated 90 degrees to view the hMPV F protein from the top down.

## Discussion

Here we demonstrate a new class of neutralizing hMPV F-specific human mAbs. The mAbs are broadly reactive across all hMPV subgroups, and neutralize viruses from both hMPV genotypes. The mAbs were discovered to bind to a novel epitope by competition with previously discovered rodent and human derived hMPV F-specific mAbs. The RSV F protein has at least two antigenic sites that are surface exposed on the head of the trimeric surface (antigenic sites Ø^60^ and V^42,43^), however, such antigenic sites have not yet been identified for hMPV F, likely due to glycan shielding^36^. Furthermore, the X-ray crystal structure of one mAb, MPV458, was determined in complex with the hMPV F protein and solved to 3.1 Å. The structure revealed MPV458 binds at a newly defined epitope on the hMPV F protein defined by the alpha helical 66-87 amino acid region contained within the F2 fragment on pre-fusion hMPV F. This new epitope is the first defined on the head of the hMPV F protein as previous mAbs identified have targeted the lower half of the protein^41,48,50–53^. The new epitope is nearly completely contained within the pre-fusion trimeric interface of the hMPV F protein, which is a unique feature among previously discovered human mAbs to viral glycoproteins. Although the mAbs were shown to bind both predominantly pre-fusion and post-fusion conformations of the hMPV F protein, preferential binding to pre-fusion hMPV F was observed as evidenced by our attempts to complex MPV458 and MPV465 with post-fusion hMPV F. These data indicate that while the 66-87 epitope is present in both pre-fusion and post-fusion conformations, the complete structural epitope is present only on pre-fusion hMPV F, as several contacts outside of the 66-87 region were observed in our X-ray crystal structure. These additional epitope residues are rearranged in the post-fusion conformation. Recently another class of human mAbs were isolated that target the influenza hemagglutinin protein^58^. The FluA-20-like mAbs were nonneutralizing, unlike the mAbs described here, which are the first human mAbs binding within the trimeric interface that neutralize a virus. As epitopes at the trimeric interface have now been determined for influenza virus^58^ and human metapneumovirus, it is likely such epitopes are important for other type I fusion viral glycoproteins.

The mechanism by which mAbs MPV458 and MPV465 neutralize hMPV remains to be determined. The mAbs could inhibit the transition of the hMPV F protein from the pre-fusion to the post-fusion conformation, which is likely the mechanism for the majority of antibodies targeting Pneumovirus fusion proteins. Alternatively, the mAbs could prevent infection by disrupting the trimeric structure of the hMPV F protein. Currently, we do not have reliable pre-fusion constructs that could be used to examine this hypothesis. It is clear that MPV458 binds to the surface of infected cells as demonstrated by our analysis by flow cytometry, although it is unclear of the mAb is binding to trimeric or monomeric hMPV F on the cell surface. Since the 66-87 epitope is hidden within the trimeric interface of the previously determined X-ray crystal structure of pre-fusion hMPV F^36^, a mechanism must occur whereby the hMPV F protein motion facilitates exposure of the epitope for MPV458 binding, and indeed for initial naïve B cell recognition of this epitope since these mAbs were derived from seropositive human subjects. This motion, termed “breathing” has previously been demonstrated for the RSV F protein by identification of an alternative conformation of the RSV F protein, whereby the mAb CR9501 causes opening of pre-fusion RSV F trimers, and RSV F was also found to be both monomeric and trimeric on the surface of transfected HEK293F cells^59^. Furthermore, breathing of influenza and HIV glycoproteins has also been described^61,62^, and mAbs to the HIV glycoprotein have been shown to destabilize the trimeric structure^63^. The mAb CR9501 targets antigenic site V of the RSV F protein, which was previously defined by the mAb hRSV90^43^. mAbs to a similar antigenic site V epitope on the hMPV F protein have not been identified, and MPV458 targets an epitope on the opposite face of monomeric hMPV F.

Although we have identified a new antigenic site by isolating two mAbs from different donors, it remains unclear if such antibodies are a major part of the hMPV F humoral immune response. It also remains to be determined if mAbs such as MPV458 will protect against viral replication *in vivo.* Since the MPV458 epitope is partially linear, as evidenced by our binding studies to reduced hMPV F, a peptide-based vaccine based solely around this epitope may elicit neutralizing antibodies. Additionally, although MPV458 and MPV465 target a similar epitope based on epitope binning analysis, the binding properties to trimeric hMPV F are quite distinct. MPV458 shows binding to both monomeric and trimeric hMPV F constructs, while binding to trimeric hMPV F is completely eliminated for MPV465. Further structural analysis of the MPV465 epitope will delineate the differential binding properties. Our findings provide novel insights on the human antibody response to the hMPV F protein, and responses to viral glycoproteins. The X-ray crystal structure of the immune complex may guide the development of vaccines against hMPV. In addition, MPV458 can be potentially applied to the treatment and prevention of hMPV infection if prophylactic efficacy is demonstrated in animal challenge models.

## Methods

### Blood draws and informed consent

This study was approved by the University of Georgia Institutional Review Board as STUDY00005127. Healthy human donors were recruited to the University of Georgia Clinical and Translational Research Unit. After obtaining informed consent, 90 mL of blood was drawn by venipuncture into 9 heparin-coated tubes, and 10 mL of blood was collected into a serum separator tube. Peripheral blood mononuclear cells (PBMCs) were isolated from human donor blood samples using Ficoll-Histopaque density gradient centrifugation, and PBMCs were frozen in the liquid nitrogen vapor phase until further use.

### Production and purification of recombinant hMPV F proteins

Plasmids encoding cDNAs for hMPV F proteins listed in **Table S1** were synthesized (GenScript) and cloned into the pcDNA3.1+ vector. The plasmids were expanded by transformation in *Escherichia coli* DH5α cells with 100 μg/mL of ampicillin (Thermo Fisher Scientific) for selection. Plasmids were purified using the EZNA plasmid maxi kit (Omega BioTek), according to the manufacturer’s protocol. To generate stable cell lines that express hMPV B2 F, hMPV B2 F-GCN4, and hMPV F 130-BV, Expi293F (Thermo Fisher Scientific) cells were plated into a 12 well plate (4 × 10^5^ per well) with 1 mL of growth medium (Dulbecco's Modified Eagle Medium (Corning), 10% fetal bovine serum (Corning)) 1 day before transfection. For each milliliter of transfection, 1 μg of plasmid DNA was mixed with 4 μg of 25,000-molecular-weight polyethylenimine (PEI; PolySciences Inc.) in 66.67 μl Opti-MEM cell culture medium (Gibco). After 30 min, the DNA-PEI mixture was added to HEK293F cells in Opti-MEM. After 3 to 4 days, 20 μl of cell culture supernatant was used for Western blot to determine protein expression. Then, the culture medium was replaced with 1 mL growth medium supplemented with G418 (Geneticin; VWR) antibiotic to a final concentration of 250 μg/mL. After 2-3 days, HEK293F cells were resuspended with the growth medium supplemented with G418, and expanded to a 25-cm^2^ cell culture flask. Cells were trypsinized once they reach 80-90% confluency and further expanded to a 75-cm^2^ cell culture flask. Again, at 80-90% confluency, trypsinized the cells were transferred to 250 mL flask in 100 mL 293 Freestyle medium (Gibco) supplemented with G418 and cultured in shaking incubator at 37°C with 5% CO_2_. For protein expression and purification, the stable cell lines were expanded in 500 mL of Freestyle293 medium supplemented with G418. The remaining constructs are expressed by transient transfection of Expi293F cells. After 5 to 7 days, the cultures were centrifuged to pellet the cells, and the supernatants were filtered through a 0.45-μm sterile filter. Recombinant proteins were purified directly from the filtered culture supernatants using HisTrap Excel columns (GE Healthcare Life Sciences). Each column was stored in 20% ethanol and washed with 5 column volumes (CV) of wash buffer (20 mM Tris pH 7.5, 500 mM NaCl, and 20 mM imidazole) before loading samples onto the column. After sample application, columns were washed with 10 CV of wash buffer. Proteins were eluted from the column with 6 CV of elution buffer (20 mM Tris pH 7.5, 500 mM NaCl, and 250 mM imidazole). Proteins were concentrated and buffer exchanged into phosphate buffered saline (PBS) using Amicon Ultra-15 centrifugal filter units with a 30-kDa cutoff (Millipore Sigma).

### Trypsinization of hMPV F

In order to generate homogeneous cleaved trimeric hMPV F, TPCK (L-1-tosylamido-2-phenylethyl chloromethyl ketone)-trypsin (Thermo Scientific) was dissolved in double-distilled water (ddH_2_O) at 2 mg/mL. Purified hMPV B2 F was incubated with 5 TAME (p-toluene-sulfonyl-L-arginine methyl ester) units/mg of TPCK-trypsin for 1 hr at 37 °C. Trimeric and monomeric hMPV B2 F proteins were purified from the digestion reaction mixture by size exclusion chromatography on a Superdex S200, 16/600 column (GE Healthcare Life Sciences) in column buffer (50 mM Tris pH 7.5, and 100 mM NaCl). Trimeric hMPV B2 F protein was identified by a shift in the elution profile from monomeric hMPV B2 F protein. The fractions containing the trimers and monomers were concentrated using 30-kDa Spin-X UF concentrators (Corning).

### Negative-stain electron microscopy analysis

All samples were purified by size exclusion chromatography on a Superdex S200, 16/600 column (GE Healthcare Life Sciences) in column buffer before they were applied on grids. Carbon-coated copper grids (Electron Microscopy Sciences) were overlaid with 5 μl of protein solutions (10 μg/mL) for 3 min. The grid was washed in water twice and then stained with 0.75% uranyl formate for 1 min. Negative-stain electron micrographs were acquired using a JEOL JEM1011 transmission electron microscope equipped with a high-contrast 2K-by-2K AMT midmount digital camera.

### Generation of hMPV F-specific hybridomas

For hybridoma generation, 10 million peripheral blood mononuclear cells purified from the blood of human donors were mixed with 8 million previously frozen and gamma irradiated NIH 3T3 cells modified to express human CD40L, human interleukin-21 (IL-21), and human BAFF^52^ in 80 mL StemCell medium A (StemCell Technologies) containing 6.3 μg/mL of CpG (phosphorothioate-modified oligodeoxynucleotide ZOEZOEZZZZZOEEZOEZZZT; Invitrogen) and 1 μg/mL of cyclosporine (Sigma). The mixture of cells was plated in four 96-well plates at 200 μl per well in StemCell medium A. After 6 days, culture supernatants were screened by ELISA for binding to recombinant hMPV B2 F protein, and cells from positive wells were electrofused as previously described.^52^ Cells from each cuvette were resuspended in 20 mL StemCell medium A containing 1× HAT (hypoxanthine-aminopterin-thymidine; Sigma-Aldrich), 0.2× HT (hypoxanthine-thymidine; Corning), and 0.3 μg/mL ouabain (Thermo Fisher Scientific) and plated at 50 μl per well in a 384-well plate. After 7 days, cells were fed with 25 μl of StemCell medium A. The supernatant of hybridomas were screened after 2 weeks for antibody production by ELISA, and cells from wells with reactive supernatants were expanded to 48-well plates for 1 week in 0.5 mL of StemCell medium E (StemCell Technologies), before being screened again by ELISA. Positive hybridomas were then subjected to single-cell fluorescence-activated sorting into 384-well plates containing 75% of StemCell medium A plus 25% of StemCell medium E. Two weeks after cell sorting, hybridomas were screened by ELISA before further expansion of wells containing hMPV F-specific hybridomas.

### Human mAb and Fab production and purification

For recombinant mAbs, plasmids encoding cDNAs for the heavy and light chain sequences of 101F,^64^ MPE8,^49^ and DS7^47^ were synthesized (GenScript), and cloned into vectors encoding human IgG1 and lambda or kappa light chain constant regions, respectively. mAbs were obtained by transfection of plasmids into Expi293F cells as described above. For hybridoma-derived mAbs, hybridoma cell lines were expanded in StemCell medium A until 80% confluent in 75-cm^2^ flasks. Cells from one 75-cm^2^ cell culture flask were collected with a cell scraper and expanded to 225-cm^2^ cell culture flasks in serum-free medium (Hybridoma-SFM; Thermo Fisher Scientific). Recombinant cultures from transfection were stopped after 5 to 7 days, hybridoma cultures were stopped after 30 days. Culture supernatants from both approaches were filtered using 0.45 μm filters to remove cell debris. mAbs were purified directly from culture supernatants using HiTrap protein G columns (GE Healthcare Life Sciences) according to the manufacturer’s protocol. To obtain Fab fragments, papain digestion was performed using the Pierce Fab preparation kit (Thermo Fisher Scientific) according to the manufacturer’s protocol. Fab fragments were purified by removing IgG and Fc contaminants using a HiTrap MabSelectSure (GE Healthcare Life Sciences) column according to the manufacturer’s protocol.

### Isotype determination for human mAbs

For determination of mAb isotypes, 96-well Immulon HB 4× ELISA plates (Thermo Fisher Scientific) were coated with 2 μg/mL of each mAb in PBS (duplicate wells for each sample). The plates were incubated at 4 °C overnight and then washed once with water. Plates were blocked with blocking buffer (2% nonfat milk, 2% goat serum in PBS with 0.05% Tween 20 (PBS-T)) and then left to incubate for 1 hr at room temperature. After incubation, the plates were washed three times with water. Isotype-specific antibodies obtained from Southern Biotech (goat anti-human kappa-alkaline phosphatase [AP] [catalog number 100244-340], goat anti-human lambda-AP [catalog number 100244-376], mouse anti-human IgG1 [Fc]-AP [catalog number 100245714], mouse anti-human IgG2 [Fc]-AP [catalog number 100245-734], mouse anti-human IgG3 [hinge]-AP [catalog number 100245-824], and mouse anti-human IgG4 [Fc]-AP [catalog number 100245-812]) were diluted 1:1,000 in blocking buffer, and 50 μl of each solution was added to the respective wells. Plates were incubated for 1 h at room temperature and then washed five times with PBS-T. The PNPP substrate was prepared at 1 mg/mL in substrate buffer (1 M Tris base, 0.5 mM MgCl_2_, pH 9.8), and 100 μl of this solution was added to each well. Plates were incubated for 1 hr at room temperature and read at 405 nm on a BioTek plate reader.

### RT-PCR for hybridoma mAb variable gamma chain and variable light chain

RNA was isolated from expanded hybridoma cells using the ENZA total RNA kit (Omega BioTek) according to the manufacturer’s protocol. A Qiagen OneStep RT-PCR kit was used for cDNA synthesis and PCR amplification. For RT-PCR, 50 μl reaction mixtures were designed with the following final concentrations: 1× Qiagen OneStep RT-PCR buffer, 400 μM deoxynucleoside triphosphate (dNTP) mix, 0.6 μM primer mix, 2 μl of Qiagen OneStep RT-PCR enzyme mix, 1 μg total of the template RNA, and RNase-free water. Three separate sets of primer mixes were used: gamma, kappa and lambda forward and reverse primers as previously described^65^. The RT-PCR was performed in a thermocycler with the following program: 30 min at 50 °C, 15 min at 95 °C, and then a 3-step cycle with 30 repeats of denaturation for 30 s at 94 °C, annealing for 30 s at 50 °C, and extension for 1 min at 72 °C, followed by 10 min of final extension at 72 °C. Samples were analyzed by agarose gel electrophoresis and purified PCR products (ENZA cycle pure kit; Omega Biotek) were cloned into the pCR2.1 vector using the Original TA cloning kit (Thermo Fisher Scientific) according to the manufacturer’s protocol. Plasmids were purified from positive DH5α colonies with ENZA plasmid DNA mini kit (Omega Biotek) and submitted to Genewiz for sequencing. Sequences were analyzed using IMGT/V-Quest.^66^ For MPV458, 2× 10^6^ of hybridoma cells were sent to GenScript for antibody variable domain sequencing.

#### Enzyme-linked immunosorbent assay for binding to hMPV F proteins

For recombinant protein capture ELISAs, 384-well plates (Greiner Bio-One) were treated with 2 μg/ml of antigen in PBS for 1 h at 37°C or overnight at 4°C. Following this, plates were washed once with water before blocking for 1 hr with 2% blocking buffer. Primary mAbs or culture supernatants were applied to wells for 1 h following three washes with water. Plates were washed with water three times before applying 25 μl secondary antibody (goat anti-human IgG Fc; Meridian Life Science) at a dilution of 1:4,000 in blocking solution. After incubation for 1 h, the plates were washed five times with PBS-T, and 25 μl of a PNPP (p-nitrophenyl phosphate) solution (1 mg/ml PNPP in 1 M Tris base) was added to each well. The plates were incubated at room temperature for 1 hr before reading the optical density at 405 nm on a BioTek plate reader. Binding assay data were analyzed in GraphPad Prism using a nonlinear regression curve fit and the log(agonist)-versus-response function to calculate the binding EC_50_ values.

### Experimental setup for biolayer interferometry

For all biosensors, an initial baseline in running buffer (PBS, 0.5% bovine serum albumin [BSA], 0.05% Tween 20, 0.04% thimerosal) was obtained. Following this, 100 μg/mL of His-tagged hMPV F protein was immobilized on anti-penta-HIS biosensor tips (FortéBio) for 120 s. For binding competition, the baseline signal was measured again for 60 s before biosensor tips were immersed into wells containing 100 μg/mL of primary antibody for 300 s. Following this, biosensors were immersed into wells containing 100 μg/mL of a second mAb for 300 s. Percent binding of the second mAb in the presence of the first mAb was determined by comparing the maximal signal of the second mAb after the first mAb was added to the maximum signal of the second mAb alone. mAbs were considered noncompeting if maximum binding of the second mAb was ≥66% of its uncompeted binding. A level of between 33% and 66% of its uncompeted binding was considered intermediate competition, and ≤33% was considered competition. For affinity studies, hMPV B2 F or hMPV F 130-BV proteins were loaded as described above, and decreasing concentrations (100/75/50/12.5/0 μg/mL) of Fabs or IgGs were analyzed for binding by association for 120 s and dissociation for 600 s. Octet data analysis software was used to analyze the data. Values for reference wells containing no antibody were subtracted from the data, and affinity values were calculating using the local and partial fit curves function. Binding curves were independently graphed in GraphPad Prism for data visualization.

### hMPV plaque neutralization assay

LLC-MK2 cells (ATCC CCL-7) were maintained in Opti-MEM (Thermo Fisher Scientific) supplemented with 2% fetal bovine serum and grown in 225-cm^2^ flask at 37 °C in a CO_2_ incubator. Two days prior to neutralization assays, cells were trypsinized and diluted in Opti-MEM at 80,000 cells/mL, 0.5 mL of cells were seeded into 24-well plates. On the day of the experiment, serially diluted mAbs isolated from hybridoma supernatants were incubated 1:1 with a suspension of infectious hMPV B2 strain TN/93-32 or hMPV A2 strain CAN/97-83 for 1 hr. Following this, cells were inoculated with 50 μl of the antibody-virus mixture for 1 hr with rocking at room temperature. Cells were then overlaid with 1 mL of 0.75% methylcellulose dissolved in Opti-MEM supplemented with 5 μg/mL trypsin-EDTA and 100 μg/mL CaCl_2_. Cells were incubated for 4 days, after which the cells were fixed with 10% neutral buffered formalin. The cell monolayers were then blocked with blocking buffer (2% nonfat milk supplemented with 2% goat serum in PBS-T) for 1 hr. The plates were washed with water, and 200 μl of mouse anti-hMPV N primary antibody (catalog number C01851M; Meridian Biosciences) diluted 1:1,000 in blocking buffer was added to each well, and the plates were incubated for 1 hr. The plates were then washed three times with water, after which 200 μl of goat anti-mouse IgG-horseradish peroxidase (HRP) secondary antibody (catalog number 5220-0286; SeraCare) diluted 1:1,000 in blocking solution was added to each well for 1 hr. Plates were then washed five times with water, and 200 μl of TrueBlue peroxidase substrate (SeraCare) was added to each well. Plates were incubated until plaques were clearly visible. Plaques were counted by hand under a stereomicroscope and compared to a virus-only control, and the data were analyzed in GraphPad Prism using a nonlinear regression curve fit and the log(inhibitor)-versus-response function to calculate the IC_50_ values.

### Western blot

Protein samples in reducing condition were mixed with loading buffer containing β-mercaptoethanol and heated at 96 °C for 10 minutes before loading on 4-12% Bis-Tris Plus gels (Invitrogen). Samples in non-reducing conditions were diluted in loading buffer without any other treatment. Samples were transferred to PVDF membranes via iBlot system (Invitrogen) and blocked with 5% blocking buffer (5% nonfat milk, 2% goat serum in PBS-T) at 4 °C overnight. Primary antibodies were diluted at 0.5 μg/mL in PBS-T and HRP-conjugated goat anti-human secondary antibody was diluted at 1:10,000 in PBS-T. Both incubations were 1 hour at room temperature with a 5x PBS-T wash in between. Substrate (Pierce ECL Western Blotting Substrate, Thermo Scientific) was added immediately before the image was taken with ChemiDoc Imaging System (BioRad).

### Crystallization and structure determination of the MPV458 Fab + B2 F complex

To generate the complex of hMPV B2 F + MPV458 Fab complex, purified trypsinized B2 F trimer was added to MPV458 Fab at a 1:2 molar ratio and incubate at 4°C overnight. To crystallize the complex, the sample was subjected to size exclusion chromatography (S200, 16/300, GE Healthcare Life Sciences) in 50 mM Tris pH 7.5, 100 mM NaCl. The fractions containing the complex were concentrated to 15 mg/mL and crystallization trials were prepared on a TTP LabTech Mosquito Robot in sitting-drop MRC-2 plates (Hampton Research) using several commercially available crystallization screens. Crystals were obtained in the Crystal Screen HT (Hampton Research) in condition F3 (0.5 M Ammonium sulfate, 0.1 M Sodium citrate tribasic dihydrate pH 5.6, 1.0 M Lithium sulfate monohydrate). Crystals were harvested and cryo-protected with 30% glycerol in the mother liquor before being flash frozen in liquid nitrogen. X-ray diffraction data were collected at the Advanced Photon Source SER-CAT beamline 21-ID-D. Data were indexed and scaled using XDS^67^. A molecular replacement solution was obtained in Phaser^68^ using the hMPV pre-fusion F structure (PDB 5WB0) and the Fab structure (PDB 4Q9Q). The structure of the complex was completed by manually building in COOT^69^ followed by subsequent rounds of manual rebuilding and refinement in Phenix^68^. The data collection and refinement statistics are shown in Table S3.

### Flow cytometry of hMPV infected LLC-MK2 cells

LLC-MK2 cells were cultured in 75-cm^2^ flask at 80-90% confluency, and then infected with hMPV (CAN/97-83) at 0.1 MOI in Opti-MEM containing 100 μg/mL CaCl2 and 5 μg/mL Trypsin-EDTA. After 48 hours, cells were washed twice with PBS and digested with Versene (Gibco) at 37 °C for 40-50 minutes. Cells were washed once with PBS then transferred to 1.5 mL tubes, pelleted and resuspended in 1 mL FACS buffer (PBS containing 5% FBS, inactivated 2% Human serum, inactivated 2% goat serum, 2 mM EDTA pH 8.0, 10% sodium azide) and incubated for 30 min to block Fc receptors. Cells were washed three times with PBS, then aliquoted in a 96 well U bottom plate for antibody staining. Mouse anti-human IgG Fc APC (BioLegend, 409306) was used for secondary antibody staining. Stained cells were fixed in 4% paraformaldehyde and data was collected with Beckman Coulter CytoFLEX flow cytometer. Data was analyzed in FlowJo.

## Acknowledgements

These studies were supported by National Institutes of Health grants 1R01AI143865 and 1K01OD026569. University of Georgia Office of the Vice President for Research, and by the National Center for Advancing Translational Sciences award number UL1TR002378.

X-ray data were collected at the Southeast Regional Collaborative Access Team (SER-CAT) 22-ID beamline at the Advanced Photon Source, Argonne National Laboratory. SER-CAT is supported by its member institutions (see www.ser-cat.org/members.html), and equipment grants (S10_RR25528 and S10_RR028976) from the National Institutes of Health. Use of the Advanced Photon Source was supported by the U.S. Department of Energy, Office of Science, Office of Basic Energy Sciences, under Contract No. W-31-109-Eng-38.

We thank Georgia Electron Microscopy at the University of Georgia for assistance with negative-stain electron microscopy, the University of Georgia Clinical and Translational Research Unit for assistance with donor identification and blood draws, and the University of Georgia Center for Tropical and Emerging Global Diseases flow cytometry core for assistance with cell sorting. This work was supported by the National Center for Advancing Translational Sciences of the National Institutes of Health under Award Number UL1TR002378. The content is solely the responsibility of the authors and does not necessarily represent the official views of the National Institutes of Health

The structure factors and structure coordinates were deposited to the Protein Data Bank under accession code XXXX.

## Notes

### Competing Interest Statement

The authors have declared no competing interest.

