## Supplemental Material for "Antibody recognition of the Pneumovirus fusion protein trimer interface"

**Supplementary Information**

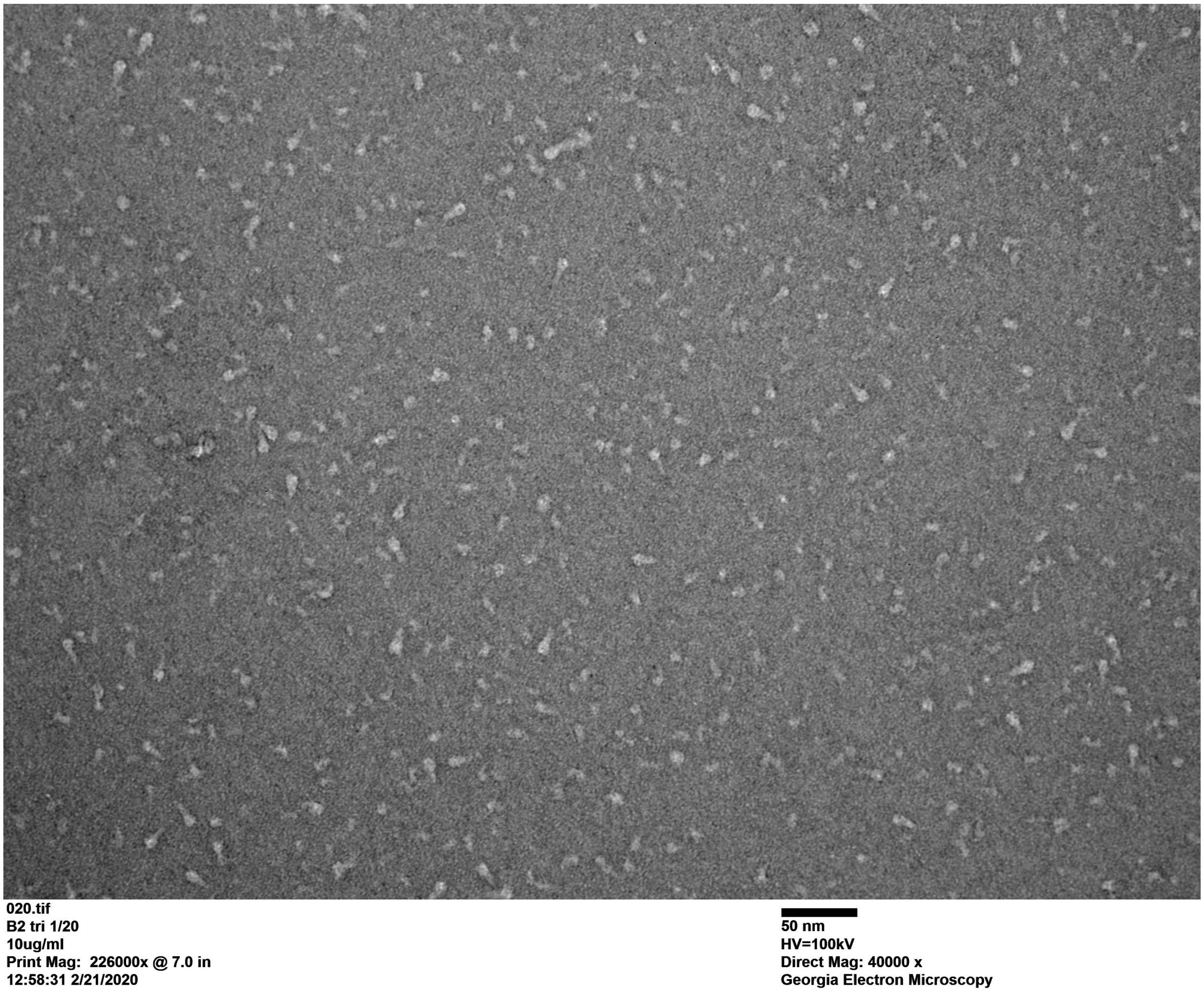


**Figure S1. Negative-stain electron micrograph of purified hMPV B2 F prior to treatment with trypsin.** A mixture of pre-fusion trimers, post-fusion trimers, and monomeric protein was observed.

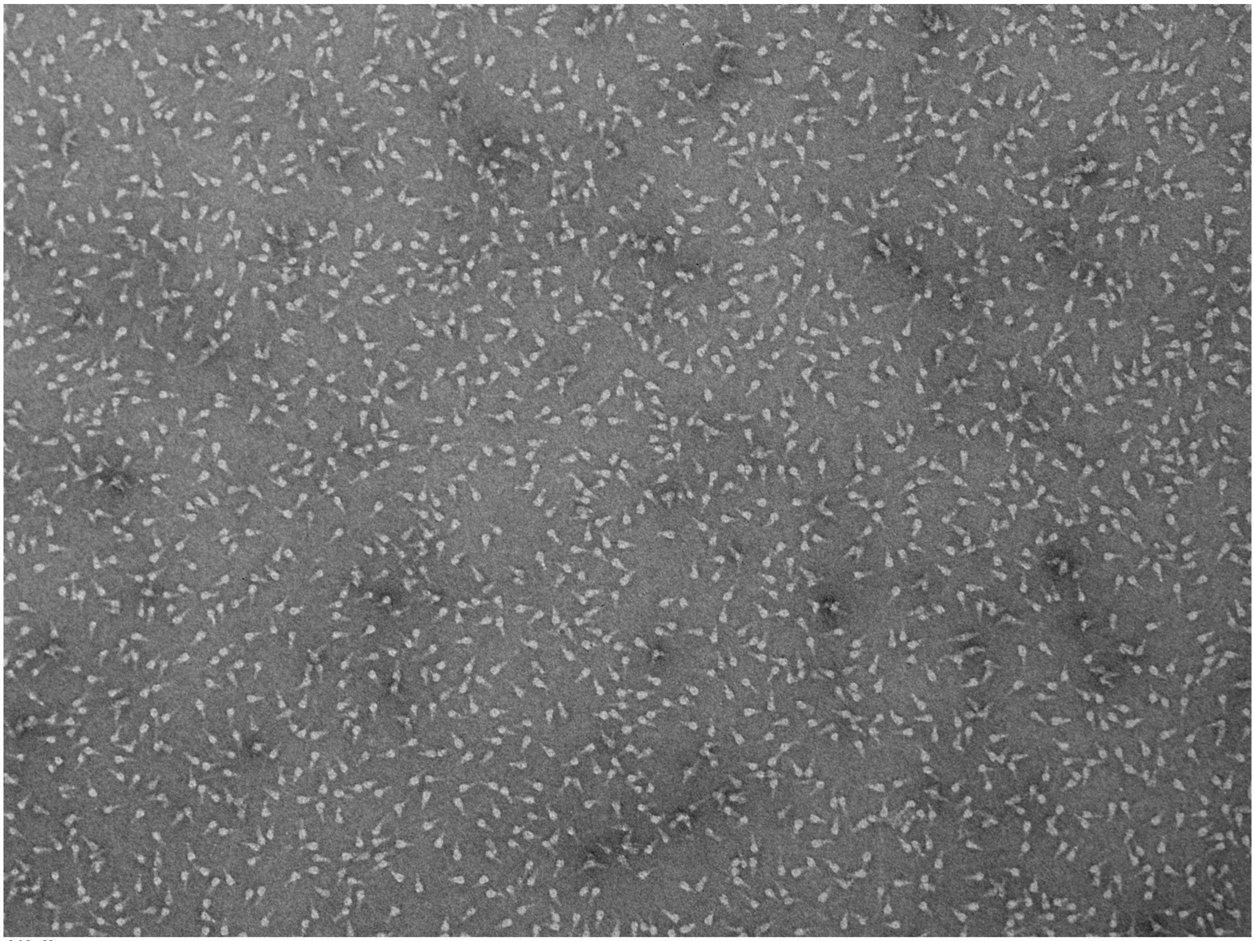

010.tif
B2opost trypsinized
10ug
Print Mag: 170000x @ 7.0 in
16:03:44 10/3/2019

100 nm
HV=100kV
Direct Mag: 30000 x
Georgia Electron Microscopy

**Figure S2. Negative-stain electron micrograph of hMPV B2 F after treatment with trypsin.** Trimeric protein was purified by size exclusion chromatography before being subjected to negative-stain electron microscopy.

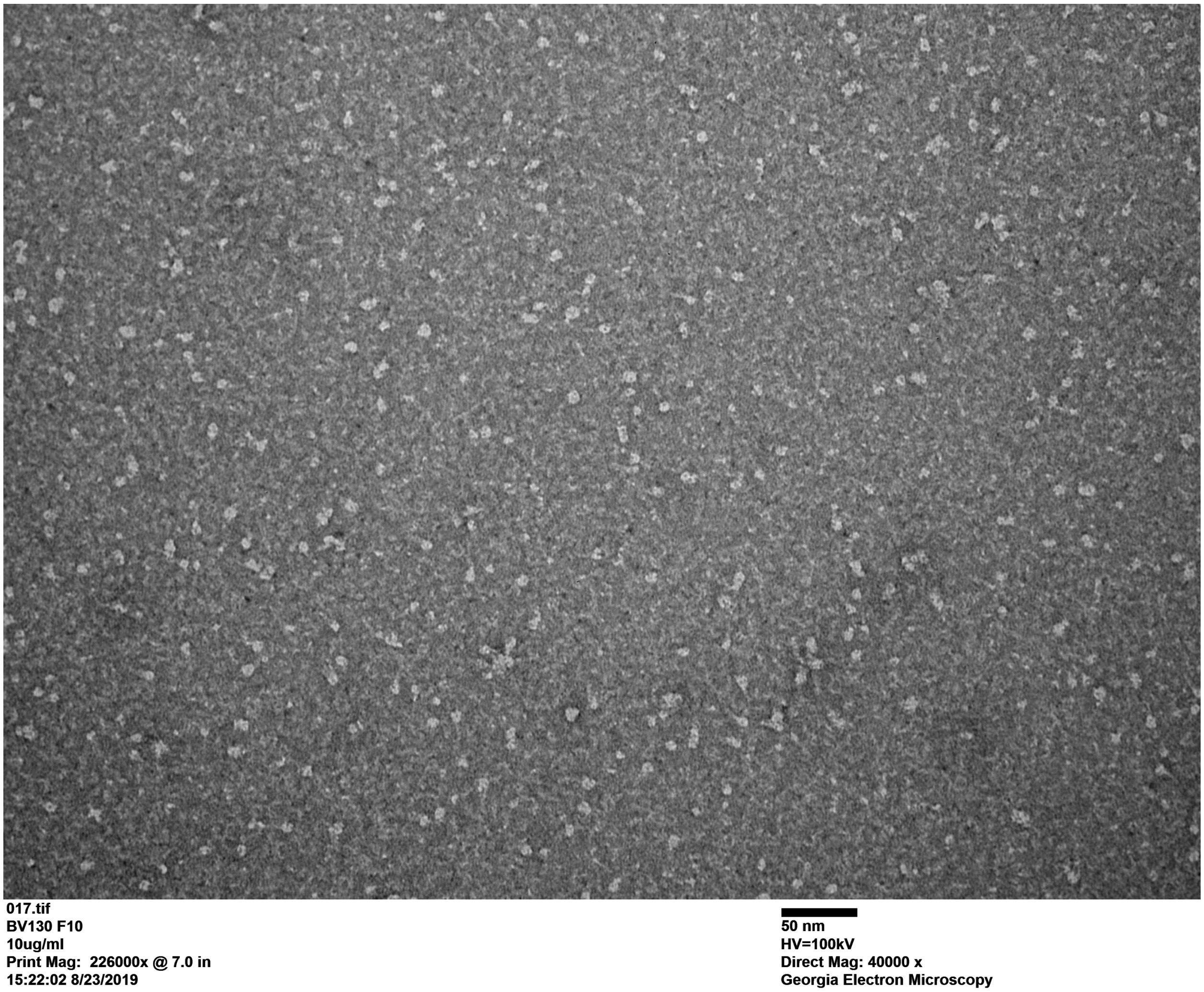


**Figure S3. Negative-strain electron micrograph of the hMPV 130-BV F protein.** The protein predominantly resembles the pre-fusion F conformation.

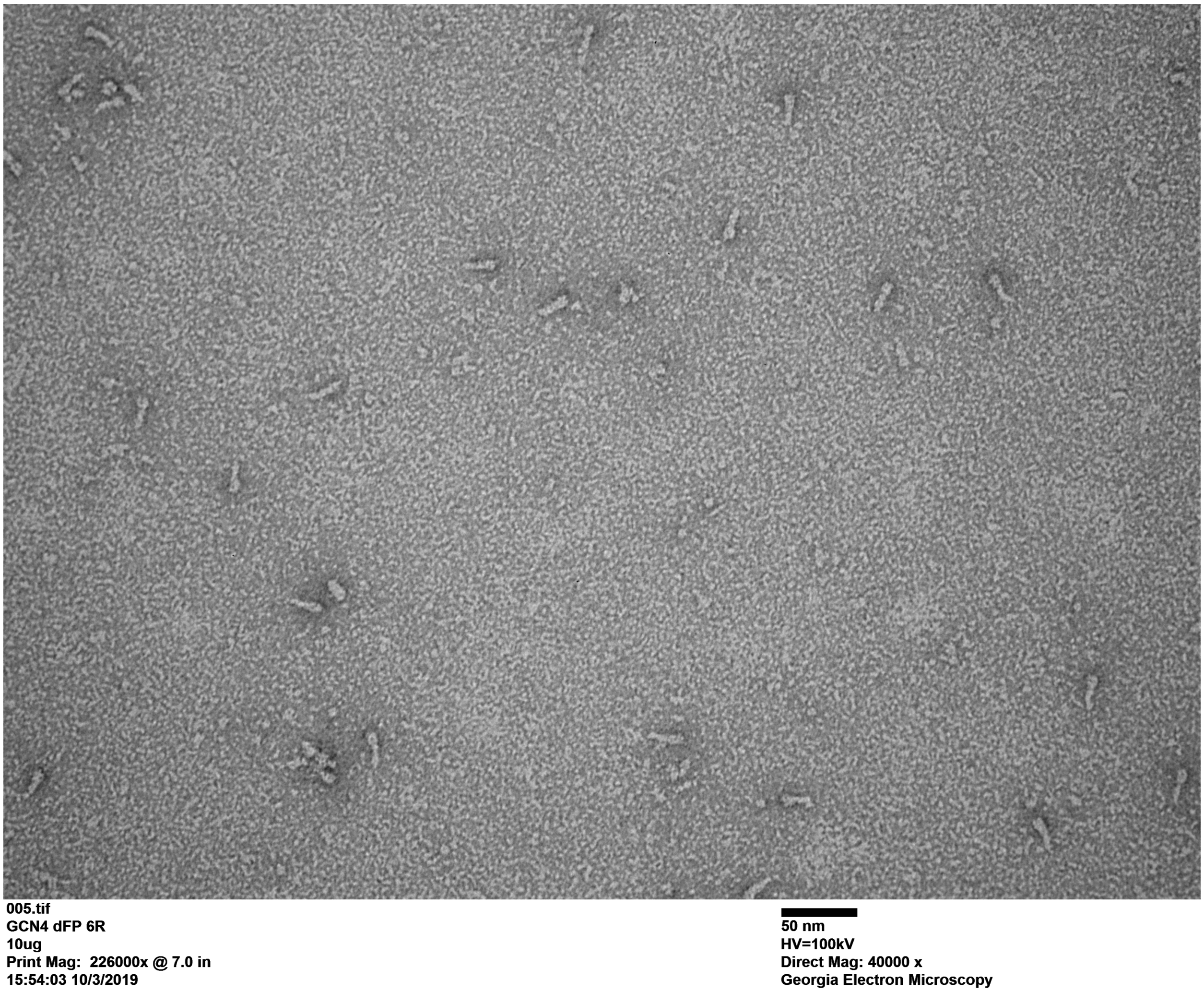


**Figure S4. Negative-strain electron micrograph of the hMPV B2 F-GCN4 dFP 6R protein.** The protein contains an extended cleavage site incorporating six repeating Arg residues to enhance intracellular cleavage. The protein predominantly resembles the post-fusion conformation.

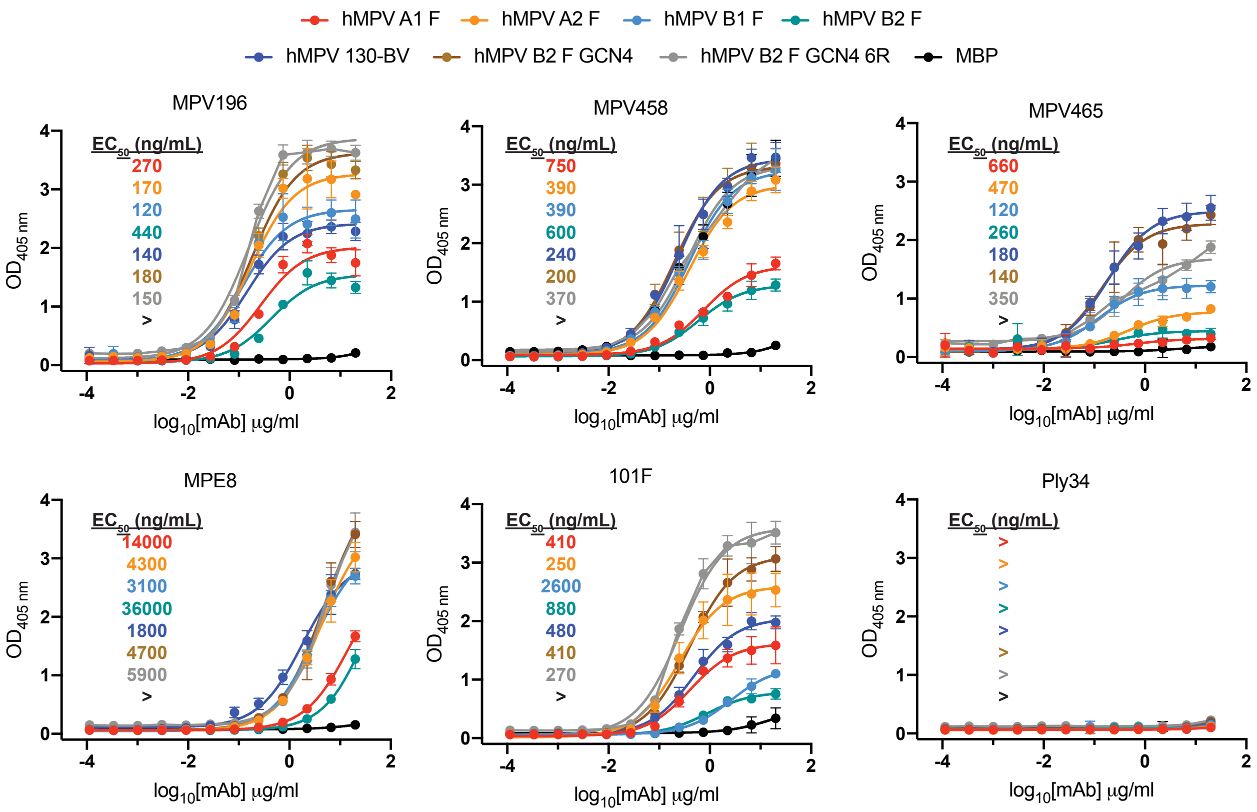


**Figure S5. ELISA binding curves of the hMPV F protein-specific mAbs to a panel of hMPV F protein constructs from multiple subgroups.** MPV458 and MPV465 bind to hMPV F proteins from all four subgroups. $EC_{50}$ values are inlaid in each graph. Binding curves and $EC_{50}$ values are colored according to the legend. Data points are the average of four replicates and error bars are 95% confidence intervals. > indicates signal above 0.5 absorbance units was not detected at the highest concentration of 20 µg/mL.

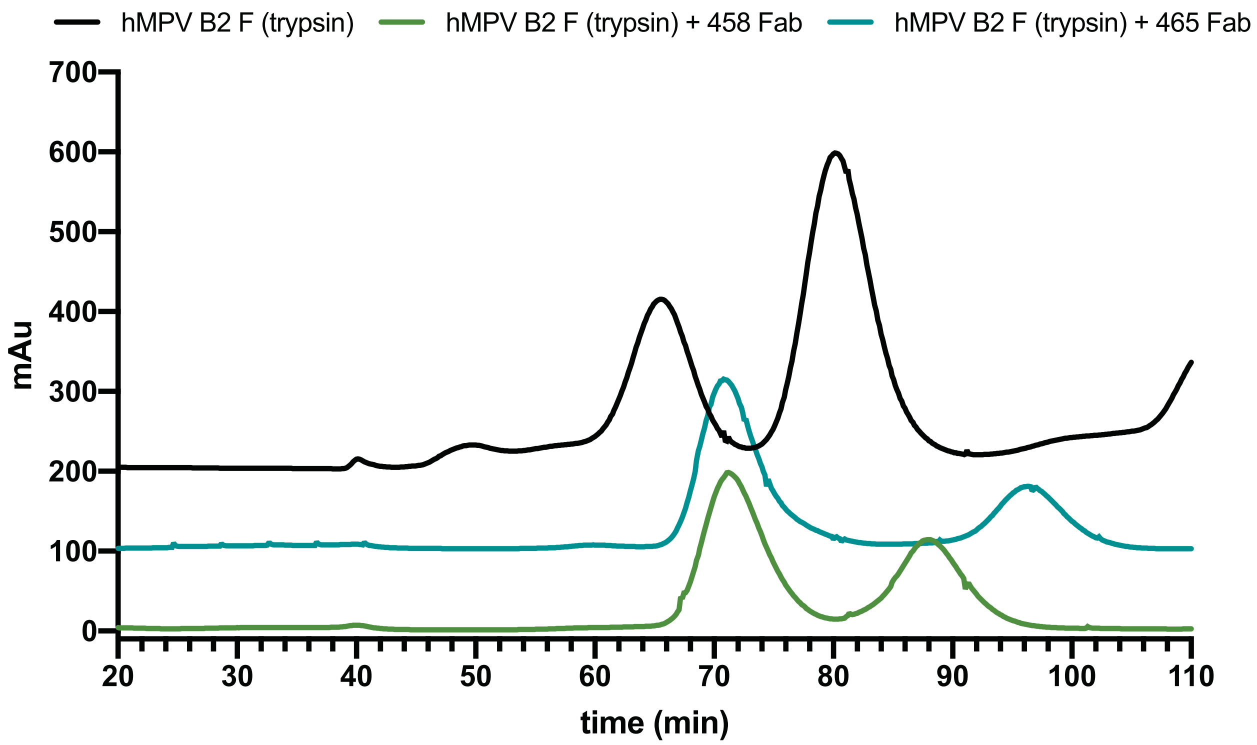


**Figure S6. Chromatograms of size exclusion chromatography of hMPV B2 F and hMPV B2 F + Fab complexes.** Trypsinization of hMPV B2 F generates homogeneous trimeric and monomeric peaks. Complexing trimeric hMPV B2 F with Fabs of MPV458 or MPV465 generates monomeric F-Fab complexes and excess Fabs. Data are representative of at least two independent experiments.

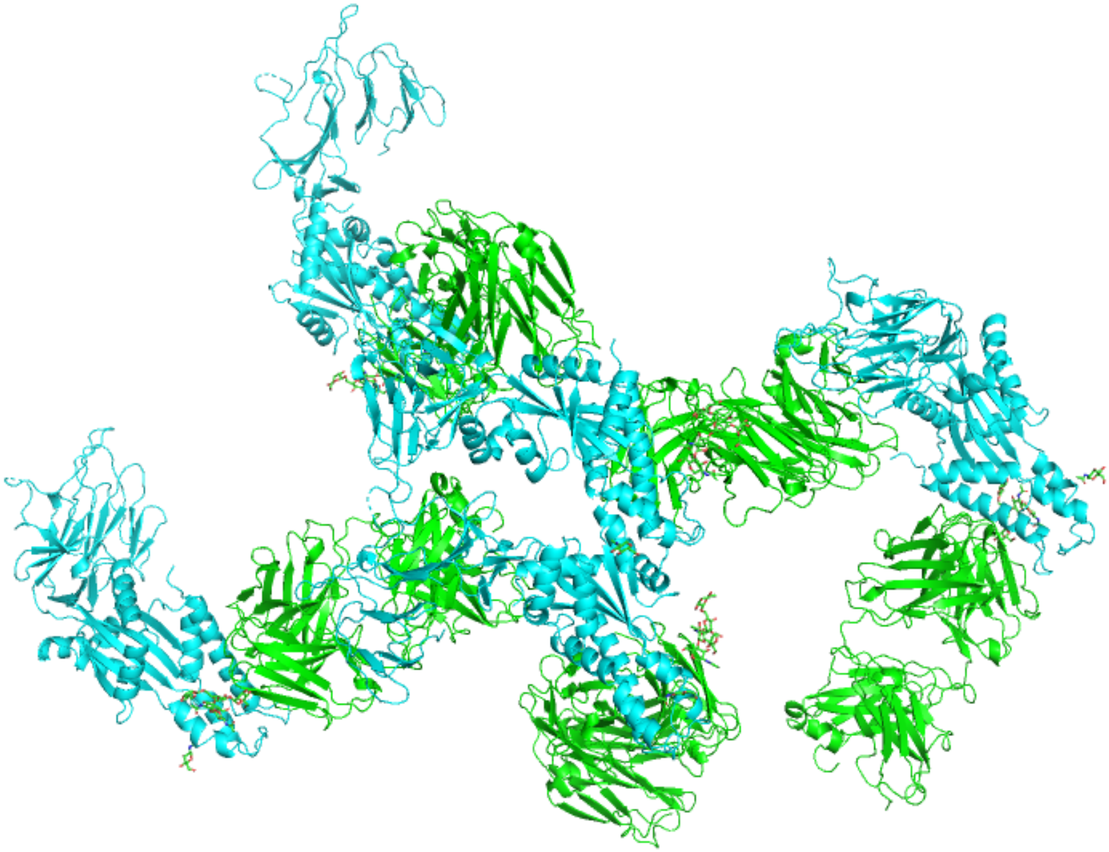

**Figure S7. Symmetry-related partners in the hMPV B2 F + MPV458 Fab complex. No trimeric structure was observed for the hMPV F protein.** The hMPV F protein is shown in cyan, while the MPV458 Fab is shown in green.

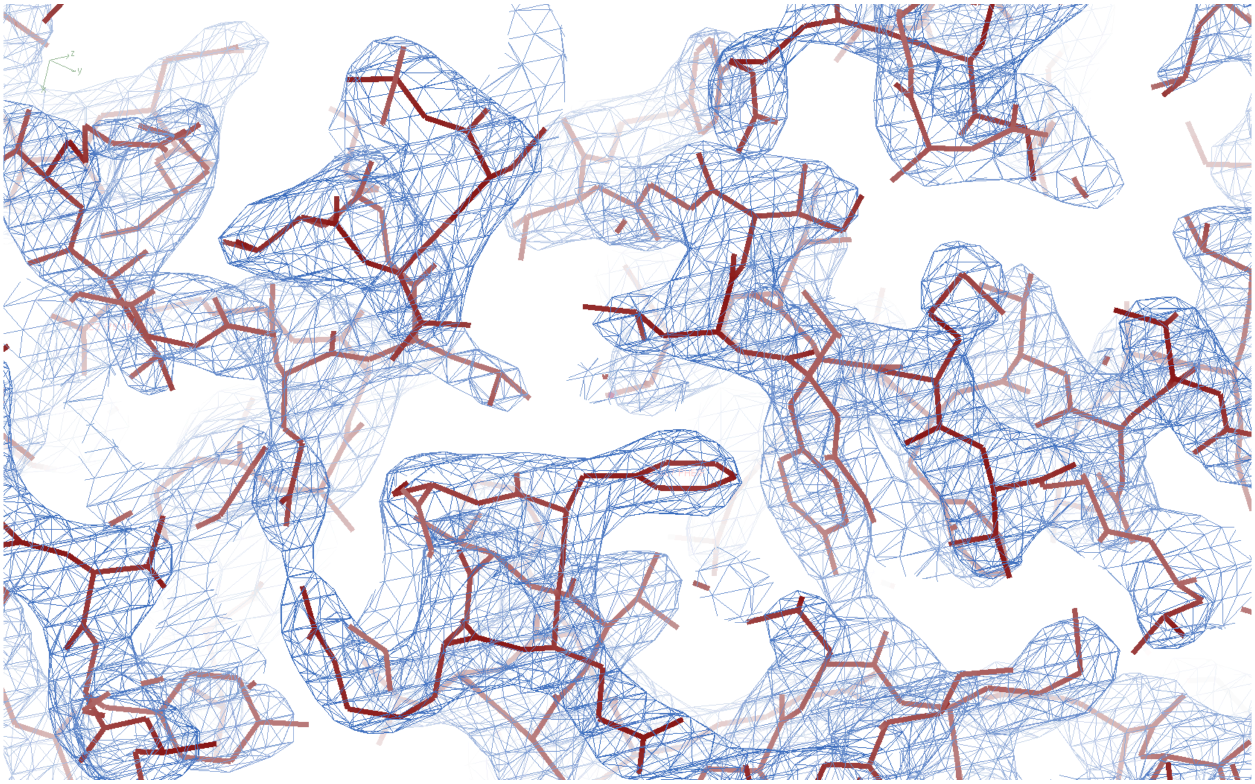

**Figure S8. Representative image of the electron density map surrounding the hMPV B2 F + MPV458 X-ray crystal structure.** The image was made in COOT and the map level rmsd is set to 2.45.

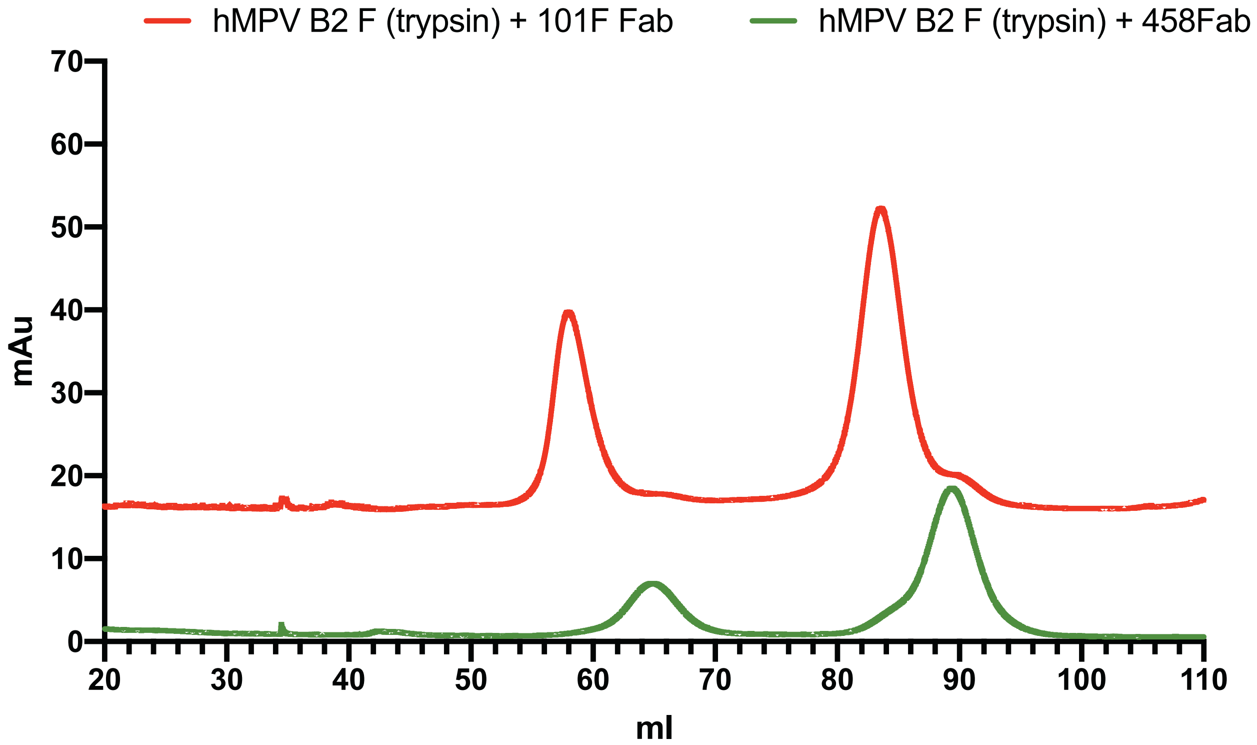


**Figure S9. Chromatographs from size exclusion chromatography of post-fusion hMPV B2 F in complex with 101F and MPV458.** No complexes were observed with MPV458, while 101F readily formed complexes with hMPV F.

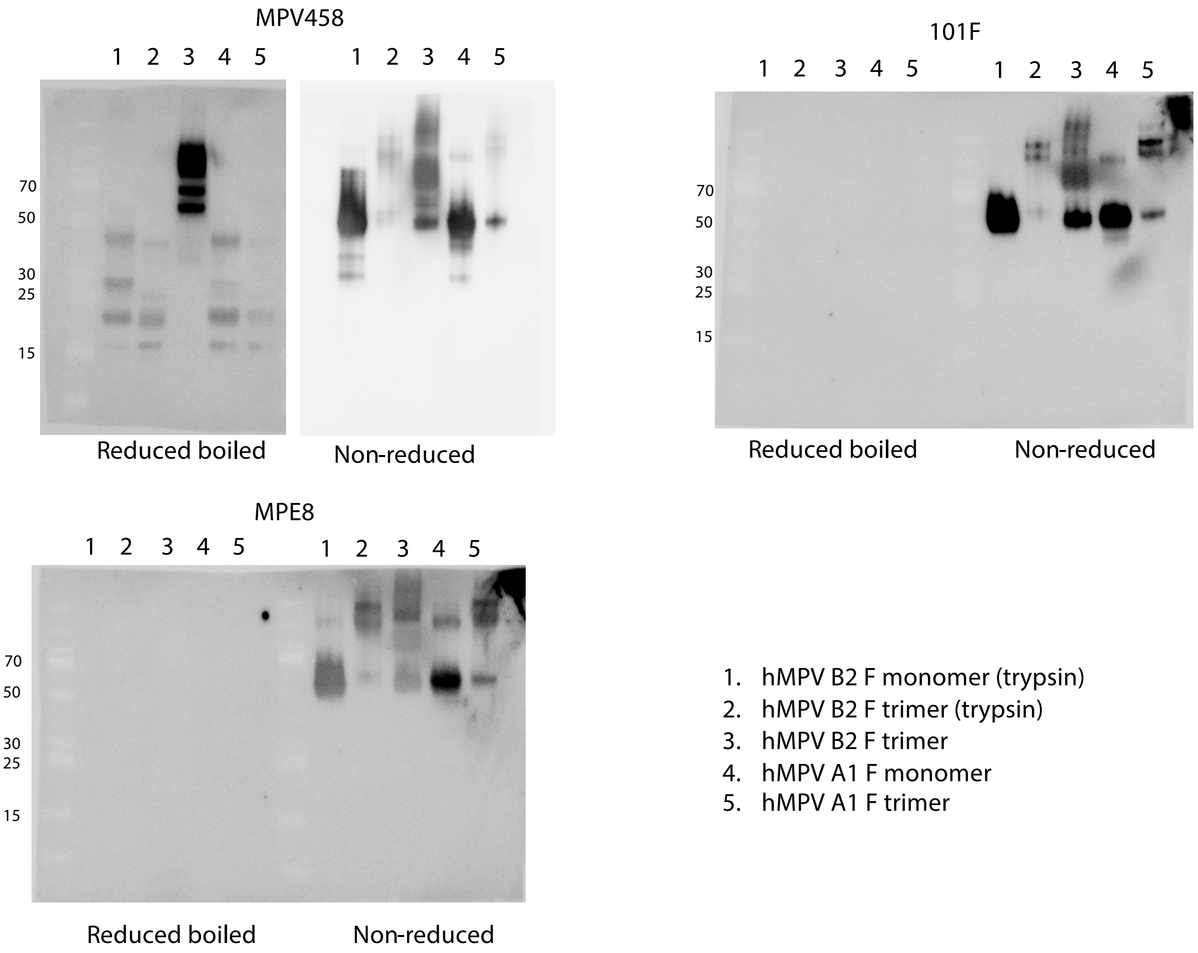


**Figure S10. Western blot analysis of MPV458 binding to the 66-87 epitope.** A panel of hMPV F protein constructs were subjected to SDS-PAGE separation before transfer to a PVDF membrane. Specific mAbs listed above each panel were used as primary antibodies. MPV458 bound to all constructs tested including boiled samples, while MPE8 and 101F bound only to samples with limited treatment.

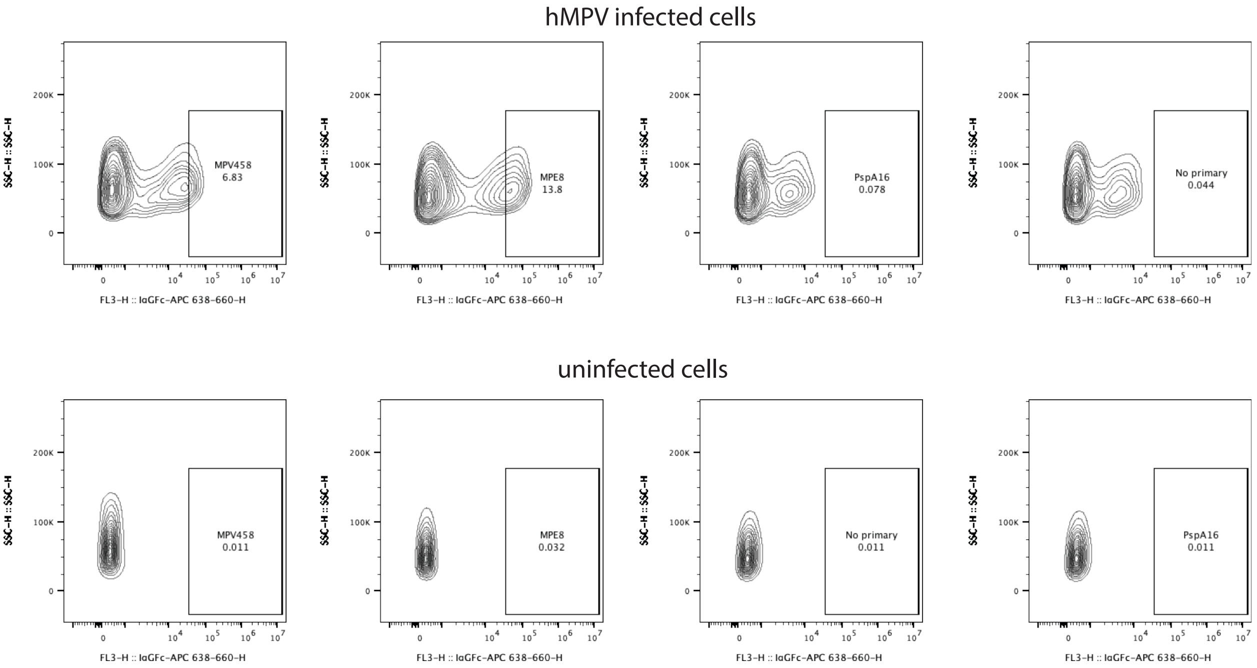


**Figure S11. Flow cytometric analysis of hMPV F infected LLC-MK2 cells.** Twenty-four hours after infection, cells were harvested and stained with mAbs indicated. MPV458 and MPE8 induced a fluorescent shift in infected cells as compared to the pneumococcal-specific mAb PspA16.

**Table S1. Properties of hMPV F recombinant protein constructs used in this study.**

| Construct name | Subgroup | Strain | Size<br>Major conformation | Cleavage site | Trimerization domain |
|---|---|---|---|---|---|
| hMPV A1 F | A1 | NL/1/00 | Monomer | KKRKRR | Foldon |
| hMPV A2 F | A2 | NL/17/00 | Monomer | KKRKRR | Foldon |
| hMPV B1 F | B1 | NL/1/99 | Monomer | KKRKRR | Foldon |
| hMPV B2 F | B2 | NL/1/94 | Monomer | KKRKRR | Foldon |
| hMPV 130-BV | A1 | NL/1/00 | Trimer<br>Pre-fusion | RQSR | Foldon |
| hMPV B2 F-GCN4 | B2 | TN/99-419 | Trimer<br>Pre-fusion | RQSR | GCN4 |
| hMPV B2 F-GCN4 dFP 6R | B2 | TN/99-419 | Trimer<br>Post-fusion | RRRRRR | GCN4 |

**Table S2. IMGT V-QUEST Analysis of MPV458 and MPV465.**

| Gene | MPV458 IgG3/kappa | MPV465 IgG1/lambda |
|---|---|---|
| **$V_H$**<br><br><br>**% identity** | IGHV3-30*03<br>IGHV3-30*18<br>IGHV3-30-5*01<br>89.6% | IGHV3-33*01<br>IGHV3-33*06<br>IGHV3-33*07<br>90.3% |
| **$D_H$** | IGHD2-2*01 | IGHD3-22*01 |
| **$J_H$**<br><br>**% identity** | IGHJ3*01<br>IGHJ3*02<br>87.8% | IGHJ5*02 F<br><br>90.2% |
| **$V_L$** | IGKV1-33*01<br>IGKV1D-33*01<br>94.3% | IGLV1-47*02 F<br><br>95.6% |
| **$J_L$** | IGKJ5*01 F<br>89.5% | IGLJ3*02<br>97.1% |
| **CDR-H1** | GFDFSRYG | GFTFGTYG |
| **CDR-H2** | IVYAGSNK | IWLDGSKT |
| **CDR-H3** | ARDQAFDL | ARAPGSVWYDTRGHMKGWFDP |
| **CDR-L1** | QGISRS | SSNIENNY |
| **CDR-L2** | DAS | GDN |
| **CDR-L3** | QQYDNLRIS | ATWDDNLSGPV |

**Table S3. Data collection and refinement statistics.**

| | hMPV B2 F + MPV458 Fab |
|---|---|
| **Wavelength** | 1.000 Å |
| **Resolution range** | 41.05 - 3.1 (3.211 - 3.1) |
| **Space group** | P 65 |
| **Unit cell** | 128.489 128.489 188.352 90 90 120 |
| **Total reflections** | 671484 (67401) |
| **Unique reflections** | 31943 (3223) |
| **Multiplicity** | 21.0 (20.9) |
| **Completeness (%)** | 99.91 (100.00) |
| **Mean I/sigma(I)** | 17.62 (1.58) |
| **Wilson B-factor** | 107.39 |
| **R-merge** | 0.1407 (2.188) |
| **R-meas** | 0.1443 (2.243) |
| **R-pim** | 0.03155 (0.4889) |
| **CC1/2** | 0.999 (0.621) |
| **CC*** | 1 (0.875) |
| **Reflections used in refinement** | 31931 (3223) |
| **Reflections used for R-free** | 3241 (350) |
| **R-work** | 0.1897 (0.3470) |
| **R-free** | 0.2339 (0.4018) |
| **CC(work)** | 0.960 (0.680) |
| **CC(free)** | 0.958 (0.511) |
| **Number of non-hydrogen atoms** | 6265 |
| **macromolecules** | 6190 |
| **ligands** | 75 |
| **Protein residues** | 818 |
| **RMS(bonds)** | 0.011 |
| **RMS(angles)** | 1.26 |
| **Ramachandran favored (%)** | 92.82 |
| **Ramachandran allowed (%)** | 6.93 |
| **Ramachandran outliers (%)** | 0.25 |
| **Rotamer outliers (%)** | 0.00 |
| **Clashscore** | 11.10 |
| **Average B-factor** | 100.80 |
| **macromolecules** | 100.36 |
| **ligands** | 137.75 |

Statistics for the highest-resolution shell are shown in parentheses.
